# DNA-terminus-dependent transcription by T7 RNA polymerase and its C-helix mutants

**DOI:** 10.1101/2023.12.07.570323

**Authors:** Bingbing Yu, Yifan Chen, Yan Yan, Bin Zhu

## Abstract

The remarkable success of mRNA-based vaccines has underscored their potential as a novel biotechnology platform for vaccine development and therapeutic protein delivery. However, the single-subunit RNA polymerase from bacteriophage T7 widely used for in vitro transcription is well known to generate double-stranded RNA (dsRNA) byproducts that strongly stimulate the mammalian innate immune response. The dsRNA was reported to be originated from self-templated RNA extension or promoter-independent transcription. Here, we identified that the primary source of the full-length dsRNA during in vitro transcription is the DNA-terminus-initiated transcription by T7 RNA polymerase. Guanosines or cytosines at the end of DNA templates enhance the DNA-terminus-initiated transcription. Moreover, we found that aromatic residues located at position 47 in the C-helix interfere with the binding of T7 RNA polymerase to DNA termini, leading to a significant reduction in the production of full-length dsRNA. As a result, the mRNA synthesized using the T7 RNA polymerase G47W mutant exhibits higher expression efficiency and lower immunogenicity compared to the mRNA produced using the wild-type T7 RNA polymerase.

## INTRODUCTION

The extraordinary success of mRNA-based vaccines against the COVID-19 pandemic highlighted the potential of mRNA-based biotechnology for vaccine development and therapeutic protein delivery (1–4). As an emerging class of medicines, mRNA-based therapeutics are considered safer, more efficient, and more economical compared to DNA-based therapeutics and conventional protein/peptide drugs (5,6). The only available method to produce large amounts of designed mRNA is the *in vitro* transcription (IVT) carried out by single-subunit RNA polymerases (ssRNAPs). However, the most popular ssRNAP, from bacteriophage T7, is known to generate undesired double-stranded RNA (dsRNA) byproducts that can stimulate the mammalian innate immune system (7,8). The dsRNA byproducts of T7 RNAP are mainly generated by self-template extension (9–13) or promoter-independent transcription (14,15). For therapeutic applications that require as much as a 1000-fold higher level of protein than vaccines to reach the therapeutic threshold, it is necessary to reduce the immuno-stimulatory effect of the dsRNA to a safe level (1–5).

Previous studies demonstrated that incorporation of modified nucleotides such as N1-methylpseudouridine (me^1^ψ) could reduce the immunogenicity of *in vitro* transcribed RNA (16–20). Certain modified nucleotides inhibit the generation of dsRNA byproducts in T7-RNAP-based IVT (15,21). Other efforts to reduce the production of dsRNA byproducts include applying high-salt transcription conditions with tight-binding promoter variants (22), co-tethering promoter DNA and T7 RNAP on magnetic beads (23), adding competing 3′-capture DNA (24) or chaotropic agents (2), and lowering the Mg^2+^ concentration (15,21). In addition, post-transcriptional purification techniques such as reverse-phase high-pressure liquid chromatography (HPLC) (18,25) or cellulose chromatography (26) can also reduce dsRNA contaminants. Nevertheless, the above methods would increase manufacturing costs, decrease yield, and/or introduce new contaminants.

Recent studies have focused on ssRNAP, the core component of IVT, to reduce dsRNA production. Lu et al. and Xia et al. reported that two novel ssRNAPs, from *Klebsiella* phage KP34 (27) and *Pseudomonas* phage VSW-3 (28,29), respectively, produced minimal self-templated dsRNA. Wu et al. reported that thermostable T7 RNAP mutants are able to synthesize functional mRNA with reduced immunogenicity at a high temperature (50℃) (30). Wu et al. reported that a single mutation S43Y attenuated RNA-dependent RNAP activity of T7 RNAP by weakening the RNA rebinding (31). Dousis et al. reported that an engineered T7 RNAP containing mutations G47A and 884G increased the 3′ homogeneity of transcribed RNA from 6–12% to more than 90% (32).

dsRNA contaminants in IVT products include short dsRNA and long dsRNA similar in length to the desired ssRNA transcript (referred to as the full-length dsRNA); the latter is more significant in triggering an immune response since it mimics the viral genome. In this work, we focused on the full-length dsRNA and solutions to reduce its production. We demonstrated that the full-length dsRNA is derived from the promoter-independent DNA-terminus-initiated transcription of T7 RNAP. We also identified the initiation sites of the promoter-independent transcription of T7 RNAP and found that the guanosines and cytosines at the end of DNA templates enhance the promoter-independent transcription. We carried out a tyrosine screen on residues 42–48 in the C-helix of T7 RNAP, which is important for association with the DNA template (31–34), and found that substitutions of G47 with aromatic amino acids attenuate the promoter-independent transcription significantly by weakening the non-specific binding of T7 RNAP to DNA termini. Among these T7 RNAP mutants, G47W was found to produce the least full-length dsRNA.

## MATERIALS AND METHODS

### Protein expression and purification

DNA fragments encoding T7 RNAP variants were inserted into the pQE82L vector and DNA fragments encoding WT SP6 RNAP were inserted into the pET28b vector with N-terminal His-tags. The vectors were transformed into *Escherichia. coli* BL21 (DE3), and cells were cultured in 1 L LB medium containing 100 mg/ml ampicillin (for T7-RNAP-pQE82L vector) or 50 mg/ml kanamycin (for SP6-RNAP-pET28b vector) at 37°C until the OD600 reached 1.2. Overexpression of RNAPs was induced by addition of 0.5 mM IPTG and continuous incubation at 16°C for 16 h. The cells were collected, resuspended in buffer (50 mM Tris-HCl, pH 7.5, 300 mM NaCl), and lysed by ultrasonication. Then, the RNAPs were purified with Ni-NTA-agarose columns (Qiagen) in buffer (50 mM Tris-HCl, pH 7.5, 300 mM NaCl) and a HiTrap Q HP column (cytiva) as per the manufacturer’s protocol. Finally, the concentrated proteins were dialyzed two times against dialysis buffer (100 mM NaCl, 50 mM Tris-HCl, pH 7.5, 1 mM DTT, 0.1 mM EDTA, 50% glycerol, 0.1% Triton X-100). VSW-3, KP34, and Syn5 RNAP were purified as described previously (27–29,35,36).

### Transcription assays

Sequences of the DNA templates and primers are shown in Table S1. All the DNA templates were prepared by PCR (Takara) unless otherwise noted. IVT reactions by T7, KP34 (27), SP6 (37), or VSW-3 RNAP (28) contained 40 mM Tris-HCl, pH 8.0, 15 mM MgCl_2_, 2 mM spermidine, 5 mM DTT, 0.2 µM inorganic pyrophosphatase, 1.5 U/µl RNase inhibitor, 4 mM each of ATP, CTP, GTP, and UTP, 30 ng/µl DNA templates, and 0.2 µM RNAP. The reaction mixtures were incubated at 37°C for 2 h (unless otherwise indicated) or at 25°C for 16 h (only for VSW-3 RNAP). IVT reactions by Syn5 RNAP containing 40 mM Tris-HCl, pH 8.0, 15 mM MgCl_2_, 2 mM spermidine, 5 mM DTT, 0.2 µM inorganic pyrophosphatase, 1.5 U/µl RNase inhibitor, 4 mM each of ATP, CTP, GTP, and UTP, 30 ng/µl DNA templates, and 2 µM RNAP were incubated at 30°C for 4 h (35). After incubation, 1 unit of DNase I was added to the reaction mixtures to remove DNA templates. In some assays, the DNase I treatment was omitted to show the DNA templates along with the transcripts. Then, the IVT products were purified with Monarch RNA purification kits (New England Biolabs, NEB), and the purified RNA was quantified using a nanophotometer (IMPLEN). To show the abortive products during the transition from transcription initiation to elongation, 0.32 mM fluorescently labeled dinucleotide 6-FAM-GG was added into the IVT reactions. The gels were imaged and analyzed using a ChemiScope 6000 Imaging System (CLINX).

### RNase III/RNase I_f_ digestion

For RNase digestion, 600 ng GFP RNA was incubated with 1, 2, 4, or 8 µl RNase III (1:1000 diluted, New England Biolabs) or RNase I_f_ (1:100 diluted, New England Biolabs) in buffer 3 (100 mM NaCl, 50 mM Tris-HCl, pH 7.9, 10 mM MgCl_2_, and 1 mM DTT, New England Biolabs) or in 50 mM NaCl, 50 mM Tris-HCl, pH 7.5, 20 mM MnCl_2_, and 1 mM DTT at 37°C for 30 min. Reactions were terminated by heat inactivation at 85°C for 5 min.

### Native gel electrophoresis

To visually distinguish ssRNA and dsRNA, IVT transcripts were analyzed by 6% TBE polyacrylamide gel electrophoresis and stained by acridine orange (AO) (Sigma Aldrich). Fluorescence gel images were obtained by a ChemiScope 6000 Imaging System (CLINX). For ssRNA-specific image, the 473-nm filter was used for excitation and the 792-nm filter was used for emission. For dsRNA-specific image, the 532-nm filter was used for excitation and the 554-nm filter was used for emission. The fluorescence images were superimposed.

### Dot blot

Various concentrations of IVT transcripts were dropped onto an Immobilon TM-Ny+ Membrane (Millipore), which was dried, blocked with 5% non-fat dry milk in TBS-T buffer (50 mM Tris-HCl, pH 7.4, 150 mM NaCl, and 0.05% Tween-20), and incubated with dsRNA-specific J2 mAb (SCICONS) for 30 min at 25°C. The membrane was washed three times with TBS-T buffer and incubated with hydrogen peroxidase-conjugated donkey anti-mouse Ig (1:2000 diluted, Jackson Immunology). After washing the membrane three times, chemiluminescence detection was performed using the ECL™ Enhanced Pico Light Chemiluminescence Kit (EpiZyme) and a ChemiScope 6000 Imaging System (CLINX).

### RNA 5′RACE and 3′RACE

A 200-nt RNA linker was prepared by IVT and its triphosphate group was converted to a monophosphate group by RppH (New England Biolabs) for 3′RACE. Then, the GFP RNA was linked to the treated linker with T4 RNA ligase 1 (New England Biolabs). The obtained RNA was purified with the Monarch RNA purification kit (New England Biolabs), and reverse transcription (RT) was performed with a specific primer using Magicscript thermotolerant reverse transcriptase (MAGIGEN). Then, the cDNA was amplified by PCR with a pair of specific primers and purified with the AxyPrep PCR Cleanup kit (Axygen). The cDNA was inserted into the plasmid pUC18 and subjected to Sanger sequencing (Genecreate). For 5′RACE, the GFP RNA was treated with RppH and linked to the 200-nt RNA linker, following the same steps as those for 3′RACE. All primers used are listed in Table S1.

### DNA binding

Various dsDNA fragments (2 µM, sequences are shown in Table S1) with or without T7 promoter were incubated with various concentrations (0, 0.5, 1, and 2 µM) of WT T7 RNAP or its G47W or E45K mutant in 40 mM Tris-HCl, pH 8.0, 15 mM MgCl_2_, 2 mM spermidine, and 5 mM DTT at 37°C for 10 min. The mixtures were mixed with 5 µl RNA dye (New England Biolabs) and separated by 12% TBE PAGE. Gels were stained with ethidium bromide, and images were analyzed by ImageJ and Prism.

### Quantitative analysis

The gray value of gel bands and immuno-blot dots was quantified with ImageJ software, and the diagrams were generated by Prism.

### mRNA synthesis

First, 3.2 mM CleanCap AG (3′ OMe) (Trilink) was added to the IVT reaction by T7 RNAP or T7 RNAP-G47W to obtain 5′-capped GFP RNA. The capped RNA was purified with the Monarch RNA purification kit, and a poly(A) tail was added to the capped RNA by *E. coli* poly(A) polymerase (New England Biolabs). The final GFP mRNA was purified with the Monarch RNA purification kit.

### Cell culture and flow cytometry

293T cells were cultured in 24-well plates (NEST) in Dulbecco’s modified Eagle medium (Gibco) supplemented with 10% fetal calf serum (Gibco), 1% penicillin/streptomycin (Thermo Fisher Scientific), and 2.5 mg/ml plasmocin prophylactic (Invivogen). At about 80% confluence, the cells were transfected with 500 ng GFP mRNA encapsulated in lipofectamine2000 (Thermo Fisher Scientific) as per the manufacturer’s protocol. GFP expression levels were recorded by fluorescence microscopy at 4, 8, and 20 h after transfection. After 20 h, fluorescence intensity was quantified by flow cytometry using SH800S (SONY).

### IFN-β detection

293T cells were cultured in 6-well plates as described above. At about 80% confluence, the cells were transfected with 2000 ng GFP mRNA encapsulated in lipofectamine2000 (Thermo Fisher Scientific) as per the manufacturer’s protocol. After 20 h, the cells were collected, lysed, and centrifuged, and the IFN-β in the supernatant was detected using the Human IFN-β (Interferon Beta) ELISA Kit (Elabscience) as per the manufacturer’s protocol.

## RESULTS

### T7 RNA polymerase produces full-length dsRNA byproducts in IVT

Four common DNA templates (*gfp*, *sox7*, the *S-gene* from SARS-CoV2, and *cas9*) were transcribed by T7 RNAP. The IVT products were analyzed by native agarose gel electrophoresis (Figure 1A) and dot blot analysis (Figure 1B). The dot blot assay detected dsRNA byproducts in all the transcripts, with the highest dsRNA content in GFP transcripts (Figure 1B). Consistently, we observed an obvious band above the gel band corresponding to the desired GFP ssRNA (Figure 1A). The position of the upper band indicates a byproduct corresponding to the full-length GFP dsRNA. To confirm the identity of this byproduct, we digested the GFP transcripts with RNase I_f_ and RNase III, which specifically degrade ssRNA and dsRNA, respectively. Then, the RNase-treated transcripts were analyzed by native agarose gel electrophoresis or PAGE, followed by ethidium bromide or acridine orange dyeing, as previously reported (15). The upper band was sensitive to RNase III but not to RNase I_f_ (Figure 1C), confirming that it presents dsRNA. As expected for the full-length dsRNA, it migrated faster than the single-stranded GFP RNA in the native PAGE (Figure 1D). Acridine orange staining further distinguished the GFP ssRNA (orange) from the full-length dsRNA (green) (Figure 1D). To reveal the mechanism of full-length dsRNA formation, we focused on the GFP transcripts with the most significant full-length dsRNA content for following investigations.

**Figure 1.**
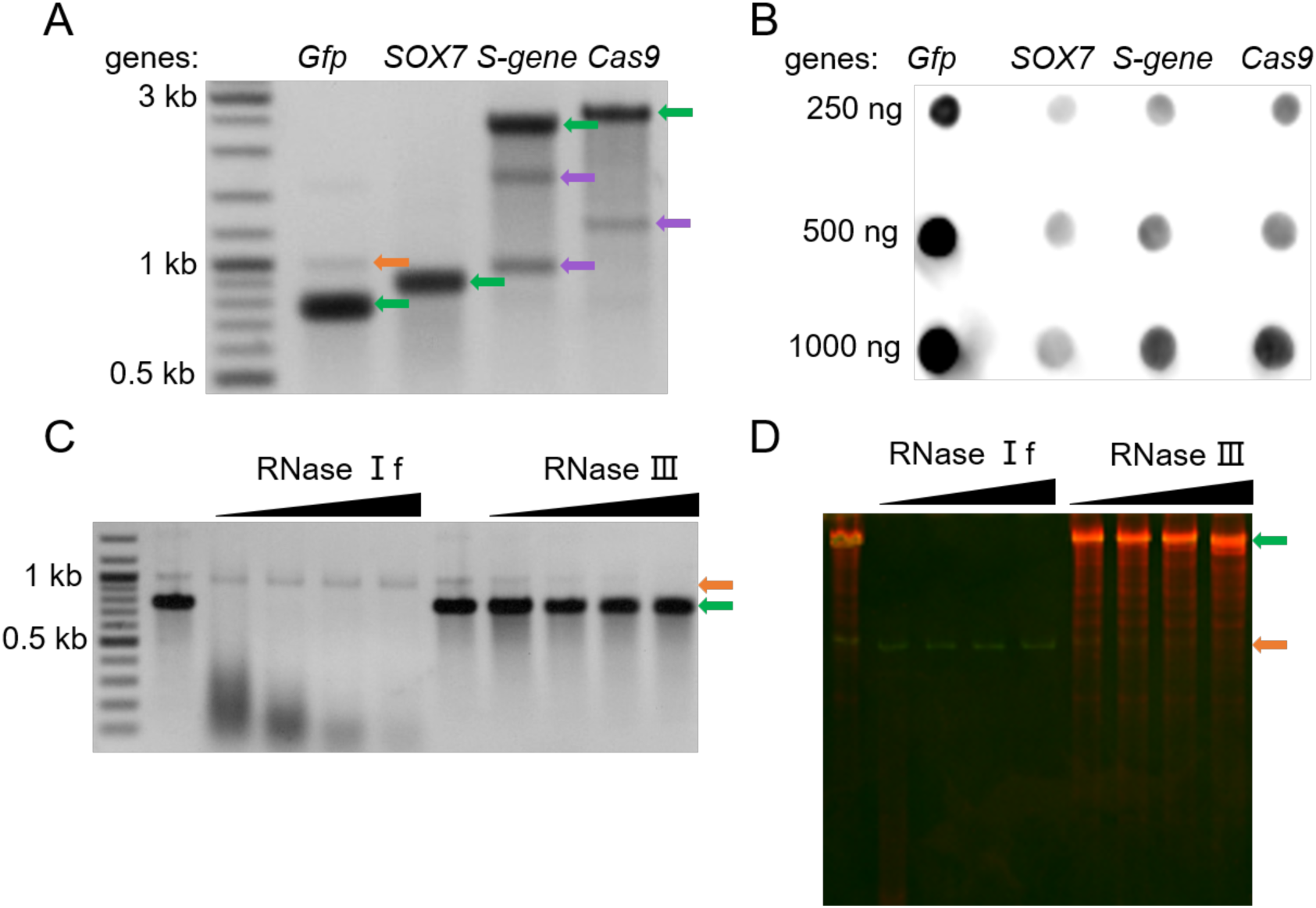
Full-length dsRNA in the IVT products of T7 RNAP. (**A**) 1.5% agarose gel electrophoresis analysis and (**B**) dot blot analysis of four genes transcribed by T7 RNAP. (**C**) 1.5% agarose gel electrophoresis analysis of GFP transcripts treated with RNase I_f_ (1/40, 1/20, 1/10, and 1/5 units/µl) and RNase III (1/10,000, 1/5000, 1/2500, and 1/1250 units/µl). (**D**) 6% native PAGE analysis of GFP tranccripts treated with RNases as in (**C**). The gel bands were visualized by acridine orange staining. Gel bands corresponding to ssRNA, dsRNA, and termintaed RNA are indicated by green, orange, and purple arrows, respectively.

### T7 RNA polymerase initiates transcription from DNA terminus without promoter

Previous studies reported that the large dsRNA byproducts of T7 RNAP were mainly generated by self-templated RNA extension (9–13) or promoter-independent transcription (15,21). To reveal the origin of the full-length dsRNA in GFP transcripts, we extended the *gfp* DNA templates with 53-, 290-, or 583-bp non-coding sequences at the 5′ termini, upstream of the T7 promoter (Figure 2A). These extensions do not affect the desired GFP ssRNA initiated from the promoter, so the size of the full-length dsRNA would not change if it originated from the self-templated RNA extension as described previously (9–13). However, if T7 RNAP initiates transcription at the 3′-termini of the DNA template, the antisense GFP transcript would be extended following the extension of the antisense DNA, and the size of the full-length dsRNA annealed by the GFP transcript (initiated at the promoter) and the antisense GFP transcript (initiated at the 3′ DNA terminus) would also increase. As shown in Figure 2A, the gel bands corresponding to the GFP ssRNA had the same mobility, despite the template 5′ extension. However, the gel mobilities of the upper bands corresponding to the full-length dsRNA were lower following the template 5′ extensions, confirming that the full-length dsRNA in GFP transcripts originated from promoter-free DNA-terminus-initiated transcription (Figure 2B) but not from RNA-templated self-extension. To further characterize the antisense transcript, we performed 3′RACE and 5′RACE analyses on the antisense RNA from the GFP DNA template with a 290-bp 5′ extension. The results of our 5′RACE analysis showed that most of the antisense RNA initiated from the second base at the non-promoter end of the DNA template (Figure 2C). 3′RACE analysis showed that the 3′ sequence of the antisense RNA matches the 5′ sequence of the antisense strand of the template DNA, confirming the run-off termination of the antisense RNA (Table S2).

**Figure 2.**
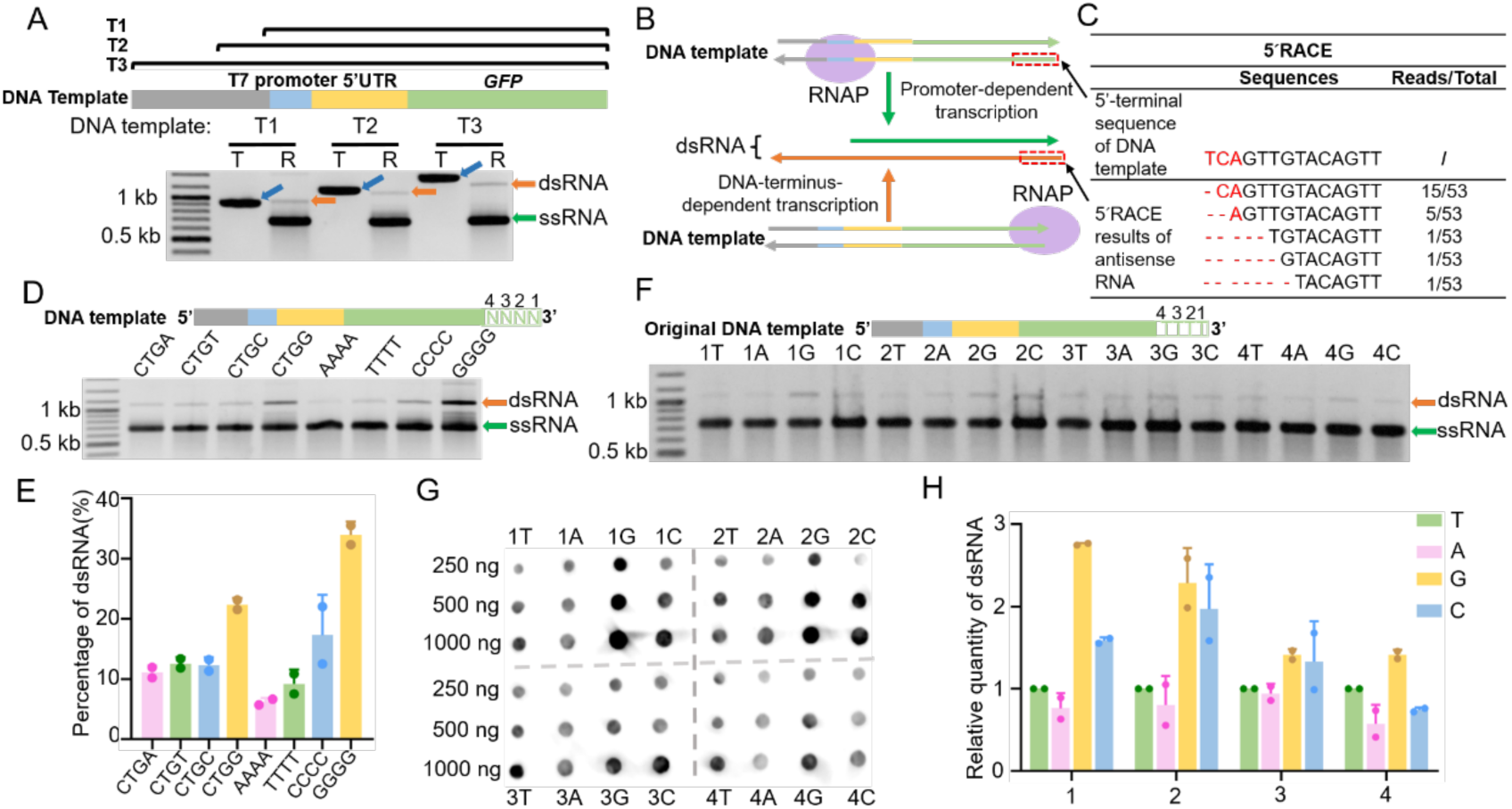
DNA-terminus-initiated transcription by T7 RNAP. (**A**) 1.5% agarose gel electrophoresis analysis of the IVT reactions by T7 RNAP on DNA templates with 5′ extensions. The non-coding sequences upstream of the T7 promoter of templates T1, T2, and T3 were 53, 290, and 583 bp in size, respectively. T indicates the DNA templates, and R indicates the transcripts. (**B**) Schematic showing the formation of full-length dsRNA in IVT. (**C**) 5′RACE results of antisense RNA. Corresponding positions of listed DNA and RNA sequences are indicated in the dashed red boxes in (**B**). (**D**) Agarose gel electrophoresis analysis of the IVT products of T7 RNAP on DNA templates with various termini. The last 4-nt sequences of DNA templates are indicated on top of the gel. (**E**) Quantification of the full-length dsRNA shown in (**D**). The gray values of the gel bands corresponding to the run-off ssRNA and the full-length dsRNA were measured, and the percentage of full-length dsRNA was calculated as the ratio between the full-length dsRNA and the sum of the run-off ssRNA and the full-length dsRNA. (**F**) Agarose gel electrophoresis analysis of the IVT products of T7 RNAP on DNA templates with various termini. The original DNA template ends with four Ts. Each of these four Ts was substituted with A, G, or C, and the production of full-length dsRNA on these templates was analyzed. (**G**) Dot blot analysis of transcripts shown in (**F**), demonstrating their full-length dsRNA contents. (**H**) Quantification of the full-length dsRNA shown in (**G**). dsRNA content was normalized to that obtained using the original DNA template (four Ts). DNA templates, ssRNA, and dsRNA are indicated by blue, green, and orange arrows, respectively, in all gels.

Previous studies (14,15) have demonstrated that the terminal structures of DNA templates influence the production of dsRNA by T7 RNAP, with the 3′ protruding DNA ends leading to more dsRNA production. The result that the production of full-length dsRNA (Figure 1A and B) varied among various DNA templates (all produced by PCR with blunt ends) indicates that the terminal sequences of DNA templates affect the DNA-terminus-initiated transcription by T7 RNAP. To clarify such influence, we prepared eight blunt-ended DNA templates with various 4-bp terminal sequences by PCR and examined their transcripts generated by T7 RNAP (Figure 2D). We calculated the proportion of dsRNA and found that DNA templates with four consecutive Gs at the ends yielded the most full-length dsRNA (Figure 2E). In contrast, DNA templates with four As or Ts at the ends yielded the least dsRNA. These results are consistent with a previous report showing that adding poly(dA) to the 3′ end of DNA templates reduces dsRNA generation in T7 RNAP IVT (30).

We further tested the effect of single variations in the 4-bp (5′-TTTT-3′) template terminal sequence on the production of full-length dsRNA. Each of the four terminal Ts was replaced by G, C, or A. The IVT products were analyzed by agarose gel electrophoresis (Figure 2F) and dot blot analysis (Figure 2G). Then we quantified the dsRNA from these templates based on Figure 2G and compared them to that from the original template (Figure 2H). Results showed that the last two bases have the most significant impact on the generation of full-length dsRNA; a single guanosine or cytidine in the terminal 2-nt region increases the yield of full-length dsRNA.

### DNA-terminus-dependent transcription is not common for bacteriophage ssRNAPs

Bacteriophages encoding ssRNAPs belong to the *Autographivirinae*, a subfamily of the *Podoviridae*, which were classified into four distinct clusters corresponding to phiKMV-, P60-, SP6-, and T7-like viruses (38) (Figure 3A). We have characterized Syn5 (36) and KP34 (27) RNAP as the representative ssRNAPs of the P60-like and phiKMV-like bacteriophages distantly related to T7, respectively. Bacteriophage VSW-3, which encodes another recently characterized ssRNAP (28), most likely also belongs to the cluster of phiKMV-like viruses based on its genomic organization (Figure 3B). We aimed to determine whether these distantly related ssRNAPs also catalyze the DNA-terminus-dependent transcription and produce the full-length dsRNA. With DNA templates harboring the same GFP coding sequence, we compared the production of the full-length dsRNA by these ssRNAPs to that by T7 and SP6 RNAP (37). Interestingly, among the five ssRNAPs investigated, KP34 and VSW-3 RNAP do not produce detectable full-length dsRNA (Figure 3C and D), indicating the ssRNAPs from phiKMV-like viruses might not initiate transcription from DNA termini. Thus, DNA-terminus-initiated, promoter-independent transcription is not a common feature for bacteriophage ssRNAPs.

**Figure 3.**
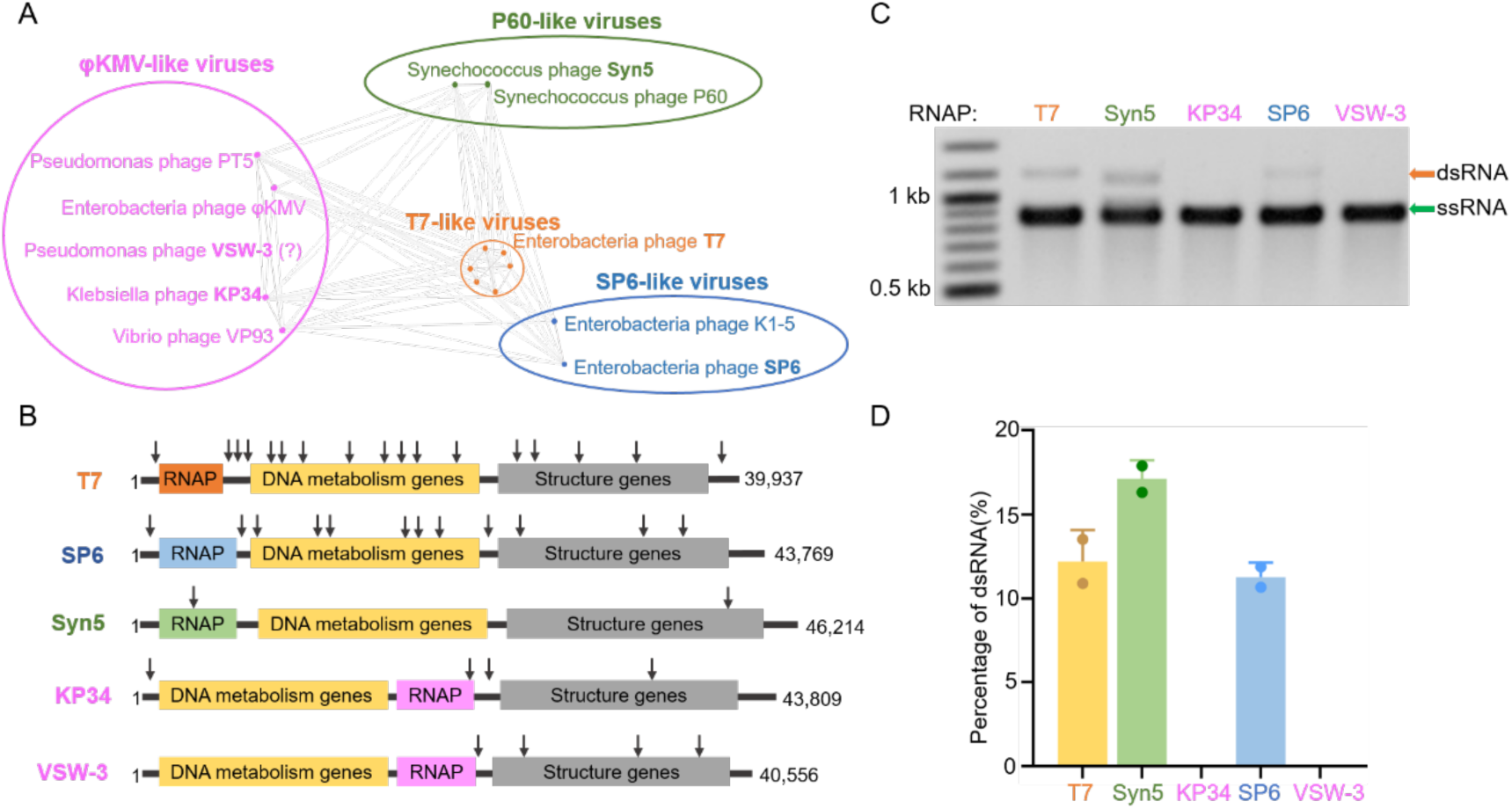
Representative ssRNAPs and their production of full-length dsRNA in IVT. **(A)** Schematic showing *Autographivirinae* clusters based on genome sequence comparison (an inaccurately modified version of Fig. 3 in (38)). **(B)** Schematic illustration of the genome organization of phage T7, SP6, Syn5, KP34, and VSW-3. The approximate positions of T7, SP6, and Syn5 promoters are indicated by downward arrows. Genome sizes are also shown. (**C**) Agarose gel electrophoresis analysis of the GFP transcripts synthesized by representative ssRNAPs in IVT. (**D**) Quantification of the results shown in (**C**). The gray values of the gel bands corresponding to the run-off ssRNA and the full-length dsRNA were measured, and the percentage of the full-length dsRNA was calculated as the ratio between the full-length dsRNA and the sum of the run-off ssRNA and the full-length dsRNA. Gel bands corresponding to ssRNA and dsRNA are indicated by green and orange arrows, respectively.

### T7 RNAP mutants with low full-length dsRNA production

Previous efforts (2,17–32) have been made to eliminate the dsRNA byproducts in IVT, which is the major source of immunogenicity for mRNA therapeutics. However, these works have barely focused on the dsRNA generated by promoter-independent antisense transcription. The fact that KP34 and VSW-3 RNAP do not initiate transcription from DNA termini encouraged us to engineer T7 RNAP to reduce its DNA-terminus-dependent transcription. Previous studies (31–34) suggested that the C-helix (residues 28–71) of T7 RNAP is important for the association between the DNA template and RNAP. The intact C-helix is formed during the transition from transcription initiation to elongation (33), with drastic conformational changes occurring at the two hinge residues, Ser43 and Gly47 (Figure 4A). Dousis et al. reported that alanine substitution at position 47 of T7 RNAP is the most advantageous for the formation of the C-helix, thus reducing the production of self-extended dsRNA to the highest extent (32). Our previous work applying directed evolution revealed that a tyrosine substitution at position 43 of T7 RNAP causes the most significant reduction of the terminal self-extended dsRNA (31). Hence, we performed a “tyrosine screen” for residues 42–48 by substituting each of the residues with tyrosine and examined the IVT products by these T7 RNAP mutants on GFP templates (Figure 4B). Interestingly, tyrosine substitutions at these positions showed various effects on the production of full-length dsRNA: E42Y and E45Y mutations increased the production of full-length dsRNA; S43Y slightly reduced the production of full-length dsRNA; while G47Y significantly reduced the production of full-length dsRNA (Figure 4C). To further confirm whether aromatic residues at position 47 cause the observed effect, we mutated G47 to Tyr, Trp, Phe, His, and Ala and analyzed their effects on the production of full-length dsRNA (Figure 4D and E). All these mutations reduced the production of full-length dsRNA by T7 RNAP, with aromatic residues showing the strongest effects (Figure 4F). Among them, the G47W mutant, with the largest residue at position 47, produced the least full-length dsRNA. The amount of full-length dsRNA in the GFP transcripts produced by the G47W mutant was reduced 9-fold compared with that produced by wild-type T7 RNAP (Figure 4F).

**Figure 4.**
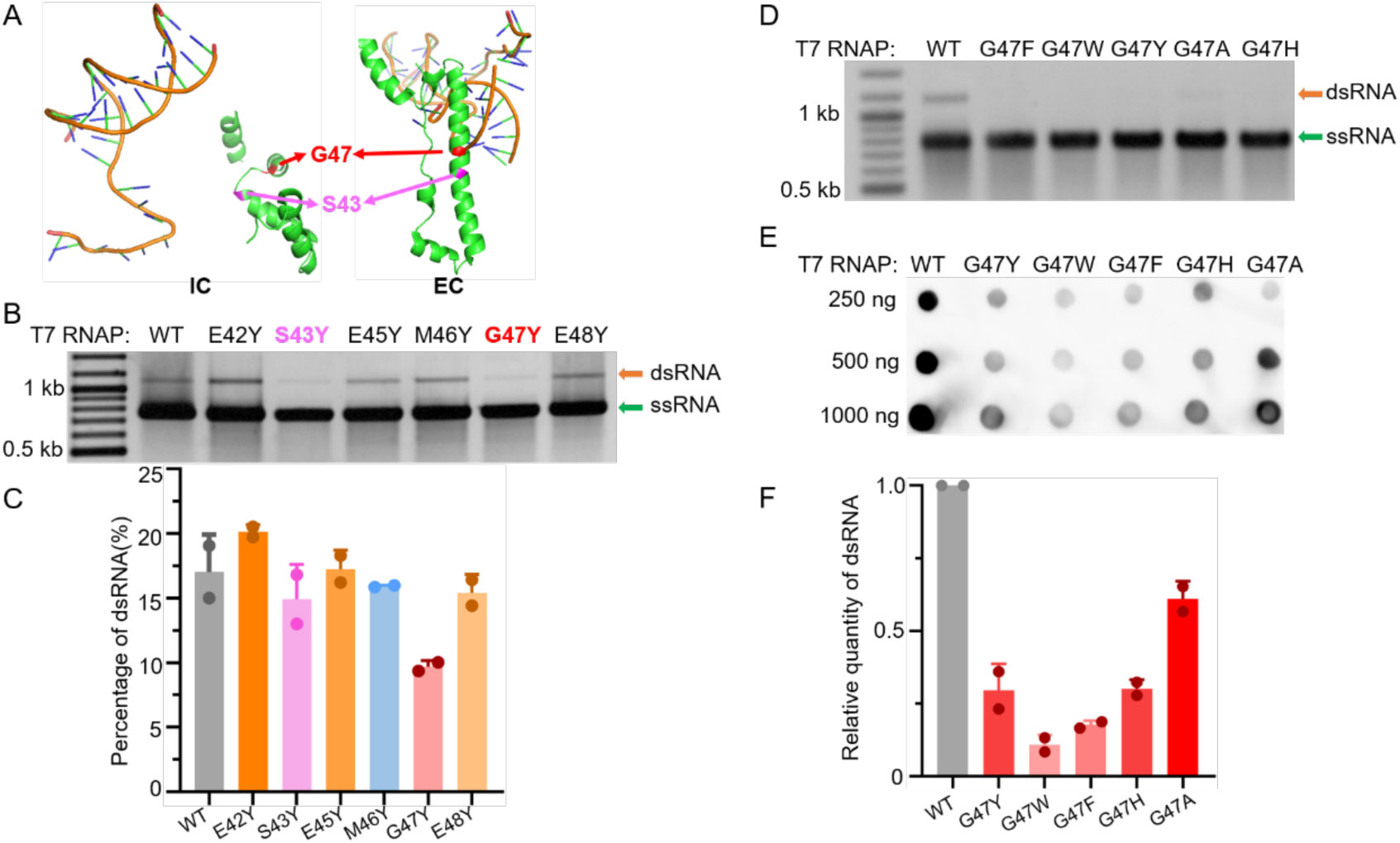
C-helix mutations in T7 RNAP affect the production of full-length dsRNA. (**A**) Structures of the C-helix in the T7 RNAP initiation complex (IC, PDB: 2PI4) and elongation complex (EC, PDB: 1MSW). Residue G47 is colored in red, and S43 is colored in purple. (**B**) Agarose gel electrophoresis analysis of GFP transcripts synthesized by wild-type T7 RNAP and its mutants E42Y, S43Y, E45Y, M46Y, G47Y, and E48Y in IVT. (**C**) Quantification of the results shown in (**B**). The gray values of the gel bands corresponding to the run-off ssRNA and the full-length dsRNA were measured, and the percentage of the full-length dsRNA was calculated as the ratio between the full-length dsRNA and the sum of the run-off ssRNA and the full-length dsRNA. (**D**) Agarose gel electrophoresis analysis of GFP transcripts synthesized by wild-type T7 RNAP and its mutants G47F, G47W, G47Y, G47H, and G47A in IVT. (**E**) Dot blot analysis of dsRNA contents from samples in (**D**). (**F**) Quantification of the full-length dsRNA shown in (**E**). dsRNA content was normalized to that obtained using wild-type T7 RNAP. Gel bands corresponding to ssRNA and dsRNA are indicated by green and orange arrows, respectively.

### G47W reduces the promoter-independent transcription by weakening the binding of T7 RNAP to DNA terminus

In contrast to G47Y, mutations E42Y and E45Y increased the production of full-length dsRNA (Figure 4C). To investigate the influence of the negative charge of these C-helix residues on the DNA-terminus-dependent transcription by T7 RNAP, we replaced the three negatively charged glutamic acids in the C-helix with positively charged lysine residues. Both gel electrophoresis (Figure 5A) and dot blot analyses (Figure 5B and C) showed significant increases in the production of full-length dsRNA in the IVT of GFP RNA by E42K, E45K, and E48K mutants compared to that by the wild-type T7 RNAP. The most significant increase was observed for the E45K mutant (Figure 5C). To elucidate the mechanism by which G47W and E45K mutations affect the DNA-terminus-dependent transcription by T7 RNAP, we designed a 40-bp DNA sequence without T7 promoter at the 5′ end but with three consecutive Gs at the 3′-terminus and investigated the binding of such DNA template by the wild-type T7 RNAP and its G47W and E45K mutants. Wild-type, G47W, or E45K T7 RNAP was incubated with the promoter-less DNA, and the binding of the enzymes to the DNA was analyzed by 12% native PAGE (Figure 5D and E). The EMSA results showed weak but specific slowly moving gel bands for wild-type T7 RNAP and the E45K mutant, indicating promoter-independent binding to the DNA terminus. However, such binding was not observed for the G47W mutant, consistent with its low dsRNA production. These results further support the notion that the full-length dsRNA originates from the promoter-independent transcription from DNA terminus.

**Figure 5.**
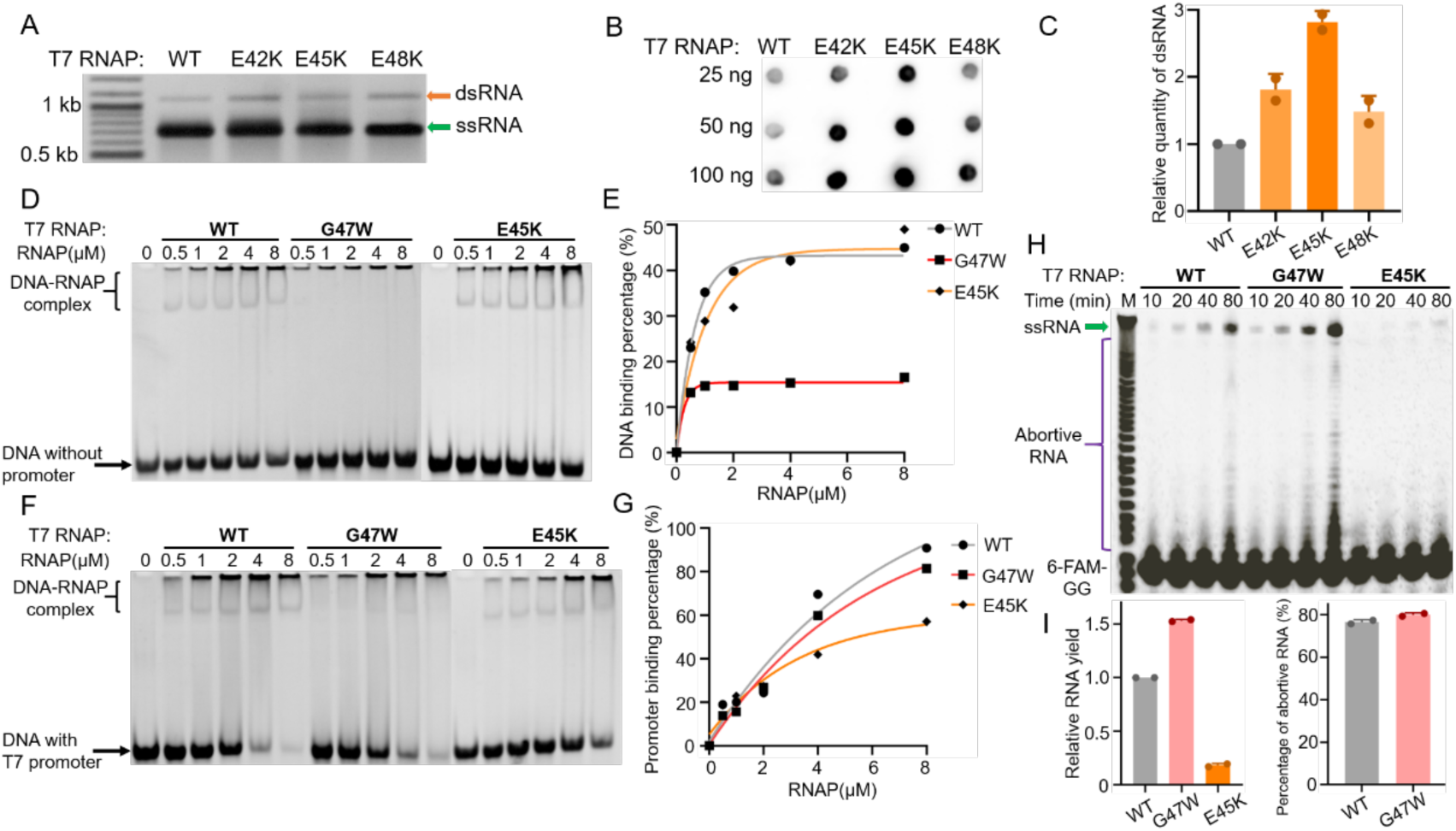
C-helix mutations affect the DNA binding by T7 RNAP. (**A**) Agarose gel electrophoresis analysis of GFP transcripts synthesized by wild-type T7 RNAP and its mutants E42K, E45K, and E48K. (**B**) Dot blot analysis of dsRNA contents from samples in (**A**). (**C**) Quantification of the full-length dsRNA shown in (**B**). dsRNA content was normalized to that obtained using wild-type T7 RNAP. (**D**) Binding of wild-type T7 RNAP or its G47W or E45K mutant to a 40-bp DNA sequence without T7 promoter. The incubated complexes were analyzed by 12% native PAGE, and the bands corresponding to unbound DNA and the DNA–RNAP complex are marked. (**E**) Quantification of the results shown in (**D**). The gray values of the gel bands corresponding to the unbound and bound DNA were measured, and the percentage of bound DNA was calculated as the ratio between the bound DNA and the sum of the bound and unbound DNA. (**F**) Binding of wild-type T7 RNAP or its G47W or E45K mutant to a 57-bp DNA sequence with a T7 promoter. The incubated complexes were analyzed by 10% native PAGE, and the bands corresponding to unbound DNA and the DNA–RNAP complex are marked. (**G**) Quantification of the results shown in (**F**). (**H**) 20% denaturing PAGE analysis of the IVT transcripts from wild-type T7 RNAP or its G47W or E45K mutant at the indicated time points. IVT reactions were initiated by the fluorescently labeled dinucleotide 6-FAM-GG, and results were visualized by fluorescence imaging. The run-off RNA, abortive RNA, and unincorporated 6-FAM-GG are marked. (**I**) Quantification of the run-off RNA shown in (**H**). The gray values of the gel bands corresponding to the run-off RNA and the abortive RNA were measured. Run-off RNA yield was normalized to that obtained using wild-type T7 RNAP in the left panel. The percentage of abortive RNA was calculated as the ratio between abortive RNA and the sum of the abortive RNA and the run-off RNA. The percentage of abortive RNA was normalized to the percentage obtained using wild-type T7 RNAP in the right panel. The abortive RNA obtained using the E45K mutant was undetectable. Gel bands corresponding to ssRNA and dsRNA are indicated by green and orange arrows, respectively.

We also investigated the effect of G47W or E45K mutation on the normal functions of T7 RNAP. First, the binding of wild-type and mutant RNAPs to the DNA containing a T7 promoter was evaluated. Inconsistent with the binding to DNA terminus, the binding to the T7 promoter was not significantly affected by the G47W mutation, while unexpectedly, the E45K mutation decreased the binding of T7 RNAP to its promoter (Figure 5F and G). These results suggest different modes of promoter or DNA terminus binding by T7 RNAP.

Moreover, as the intact C-helix is formed during the transition from transcription initiation to elongation (33), we also investigated the influence of G47W or E45K mutation on this step. T7 RNAP is known to initiate transcription efficiently with dinucleotides matching the initial RNA sequences (39,40), so we designed a 53-bp template with a T7 promoter, followed by three consecutive Gs, and added the fluorescently labeled dinucleotide 6-FAM-GG into the reactions to initiate the IVT by wild-type T7 RNAP or its G47W or E45K mutant. At 10, 20, 40, and 80 min after initiation of the reactions, 0.5 µl of every sample was taken and analyzed by 20% denaturing PAGE. Then, fluorescence gel image of RNA transcripts was obtained using the 492-nm and 517-nm filters for excitation and emission, respectively, and the image was converted into black/white (Figure 5H). We quantified the yield of run-off ssRNA and the abortive RNA products. The result demonstrated that the G47W mutant initiates transcription more efficiently with the dinucleotide GG compared to wild-type T7 RNAP to produce more run-off and abortive RNA (Figure 5I). However, the ratio between the abortive and run-off products was not significantly changed by the G47W mutation (Figure 5H and I), indicating that the transition from transcription initiation to elongation was not affected. In contrast, the E45K mutation severely reduced the initiation and the yield in IVT (Figure 5H and I), consistent with its effect on promoter binding.

### T7 RNAP-G47W is an ideal engineered RNA polymerase for mRNA production

To evaluate the advantages of T7 RNAP-G47W in mRNA production, we compared wild-type T7 RNAP and T7 RNAP-G47W in the IVT production of GFP, Cas9, and S-gene RNA. Another mutant of T7 RNAP, G47A+884G, which was recently reported to significantly reduce the self-templated dsRNA (32), was also included in the comparison. For both the G47W and G47A+884G mutants, the gel bands corresponding to the full-length dsRNA in the GFP IVT products were not observable (Figure 6A), indicating that DNA-terminus-initiated transcription by the G47A+884G mutant is also reduced compared to that by the wild-type T7 RNAP. The IVT yield of GFP RNA, with a relatively short length (873 nt), was similar for wild-type T7 RNAP and both mutants (Figure 6A and B). However, when synthesizing large RNA molecules like Cas9 (4314 nt) and S-gene (3975 nt), the IVT yield of the G47W mutant was slightly lower than that of wild-type T7 RNAP, while the IVT yield of the G47A+884G mutant was obviously reduced (Figure 6C–F). In addition, the G47A+884G mutant produced more terminated products compared to the wild-type and G47W T7 RNAP (Figure 6C and E, indicated by purple arrows).

**Figure 6.**
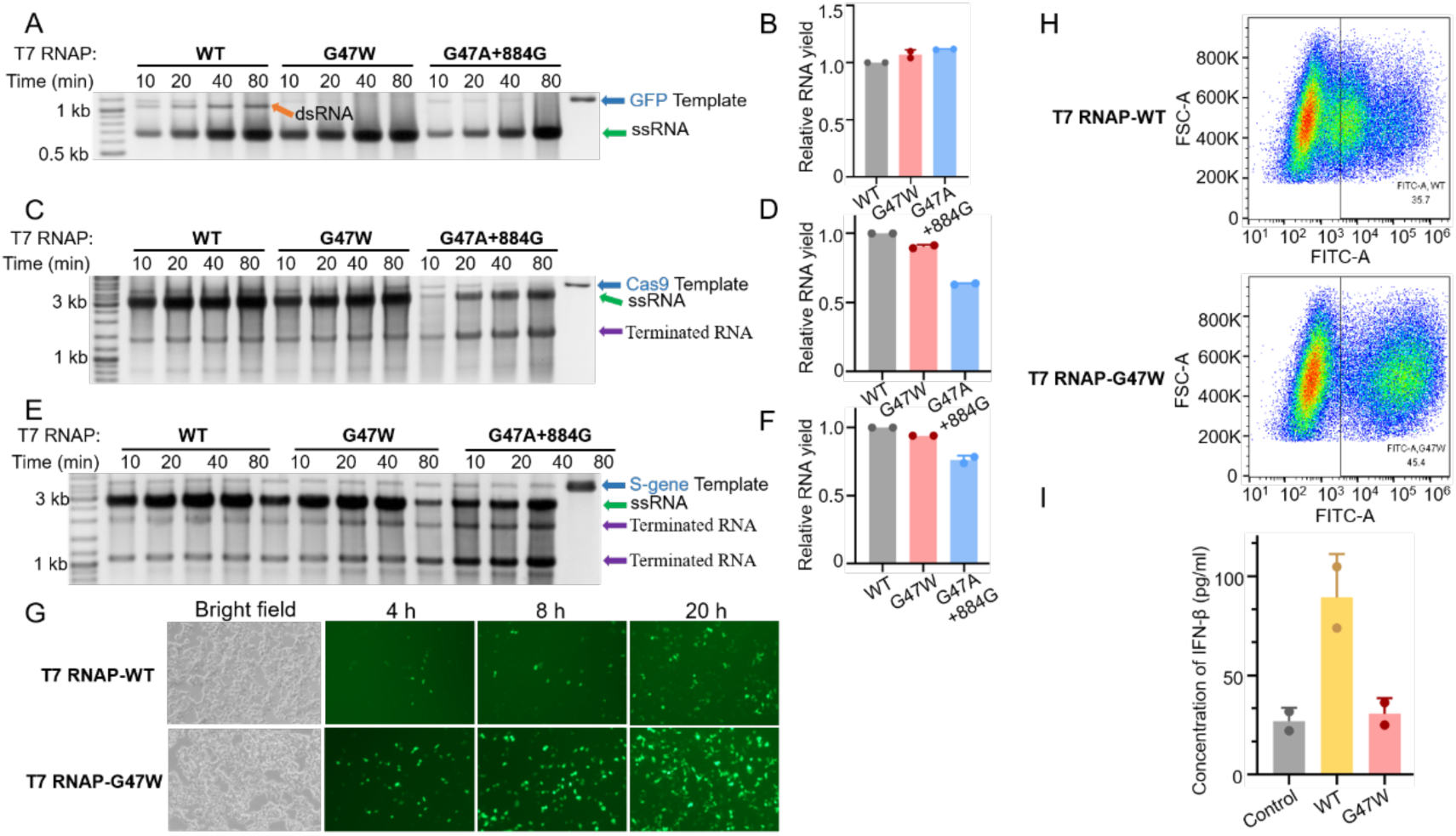
T7 RNAP-G47W is advantageous for mRNA synthesis. (**A**) 1.5% agarose gel electrophoresis analysis of the IVT production of GFP RNA by T7 RNAP (WT) or its G47W or G47A+884G mutant. (**B**) Quantification of the run-off ssRNA shown in (**A**). The gray values of the gel bands corresponding to the run-off ssRNA were measured. The run-off ssRNA yield was normalized to that obtained using wild-type T7 RNAP. (**C**) 1.5% agarose gel electrophoresis analysis of the IVT production of Cas9 RNA by T7 RNAP (WT) or its G47W or G47A+884G mutant. (**D**) Quantification of the run-off ssRNA shown in (**C**). (**E**) 1.5% agarose gel electrophoresis analysis of the IVT production of S-gene RNA by T7 RNAP (WT) or its G47W or G47A+884G mutant. (**F**) Quantification of the run-off ssRNA shown in (**E**). DNase I treatment was omitted for all reactions shown in panels **(A**), (**C**), **and** (**E**), and the DNA template is shown as a loading control. DNA templates, ssRNA, dsRNA, and terminated RNA are indicated by blue, green, orange, and purple arrows, respectively, in panels **(A**), (**C**), **and** (**E)**. (**G**) Expression levels of GFP mRNA produced by T7 RNAP-WT or T7 RNAP-G47W in 293T cells recorded by fluorescence microscopy at 4, 16, and 20 h after transfection. (**H**) Fluorescence intensity of cells transfected with GFP mRNA synthesized by T7 RNAP-WT or T7 RNAP-G47W at 20 h after transfection, as determined by flow cytometry. (**I**) Concentrations of IFN-β in cells transfected with GFP mRNA synthesized by T7 RNAP-WT or T7 RNAP-G47W at 20 h after transfection. The cells treated with lipofectamine2000 alone served as the control.

To compare the expression efficiency of mRNA produced by wild-type T7 RNAP and the G47W mutant, we transfected 293T cells with GFP mRNA synthesized by either enzyme, and their expression levels were recorded by fluorescence microscopy at 4, 16, and 20 h after transfection (Figure 6G). After 20 h, we quantified their fluorescence intensity by flow cytometry (Figure 6H). As expected, mRNA transcribed by the G47W mutant showed much higher expression levels than that transcribed by wild-type T7 RNAP. We also determined the concentration of IFN-β in 293T cells at 20 h after transfection by ELISA and found that almost no IFN-β response was induced in the cells transfected with GFP mRNA produced by the G47W mutant compared to the negative control (Figure 6I). However, the mRNA produced by wild-type T7 RNAP elicited a strong IFN-β response. These results are consistent with the low production of full-length dsRNA by the T7 RNAP-G47W mutant and demonstrate its value as an advantageous enzymatic tool for mRNA medicine and vaccine.

## DISCUSSION

It is well known that mRNA transcribed by T7 RNAP can stimulate the mammalian innate immune system and that it is necessary to reduce the immuno-stimulatory effect of the dsRNA to fit therapeutic applications. Previous studies (9–11,13) focused on the dsRNA generated by self-template extension. Recently, Ma et al. reported that wild-type T7 RNA polymerase can initiate transcription from the end of the DNA in a promoter-independent manner to generate full-length dsRNA and that this transcription could be suppressed by low concentrations of Mg^2+^ (15). Yet, the mechanisms underlying such non-conventional transcription remain elusive.

In the present study, we further analyzed the promoter-independent transcription by T7 RNAP and revealed more details about the process. We demonstrated that the production of antisense RNA is mostly initiated from the penultimate position of the DNA terminus and that the presence of guanosine or cytidine in the 2-nt terminal region of the DNA template strengthens the promoter-independent transcription by T7 RNAP significantly. Therefore, adding poly(A) to the end of DNA templates is an effective way to reduce the amounts of full-length dsRNA in mRNA production.

Moreover, we tested the full-length dsRNA production by various ssRNAPs and found that DNA-terminus-dependent transcription is not common for bacteriophage ssRNAPs. Although T7, SP6, and Syn5 RNAP as representative ssRNAPs all produce significant amounts of full-length dsRNA in IVT, the products of KP34 and VSW-3 RNAP from phiKMV-like viruses contain non-detectable dsRNA generated by promoter-independent transcription.

Previous studies proved that mutations S43Y and G47A in T7 RNAP were able to attenuate the self-template extension (31,32). In the structure of the elongation complex of T7 RNAP, this region is in close proximity to the DNA template (33,34) (Figure 4A). We replaced residues 42–48 of T7 RNAP with various amino acids, and the results showed that substitutions of G47 with aromatic amino acids most significantly reduce the dsRNA generated from promoter-independent transcription. In contrast, substitutions of negatively charged E42, E45, or E48 with lysine enhanced the promoter-independent transcription. Moreover, we demonstrated that compared to wild-type T7 RNAP, the G47W mutant showed no detectable binding to DNA terminus. These results indicate that aromatic amino acids at position 47 hinder the binding of T7 RNAP to the DNA terminus, most likely due to spatial steric hindrance. In summary, our results demonstrate the importance of residues 42–48 of T7 RNAP for the interaction with DNA; these residues may serve as potential targets for further engineering.

In summary, we generated a superior T7 RNAP mutant, G47W, with minimal dsRNA production while maintaining high yield, especially in the production of large mRNA molecules. The mRNA synthesized by T7 RNAP-G47W shows high expression efficiency and low immunogenicity, indicating that T7 RNAP-G47W is suitable for therapeutic applications.

## SUPPLEMENTARY DATA

Supplementary data are available at NAR online.

## ACKNOWLEDGEMENT

We thank all lab members for helpful discussion.

## AUTHOR CONTRIBUTION STATEMENT

B.Z. conceived the project. B.B.Y. and B.Z. designed the experiments. B.B.Y., Y.F.C. and carried out the experiments. B.B.Y., Y.F.C., Y.Y. and B.Z. analyzed the data. B.B.Y. and B.Z. wrote the manuscript. All authors discussed the results and contributed to the final manuscript.

