## Supplementary materials for "DNA-terminus-dependent transcription by T7 RNA polymerase and its C-helix mutants"

**Table S1**. Sequences of the DNA templates and primers used in this study.

| DNA | Sequence (5´-3´) |
|---|---|
| *Gfp* | ATGGCTAGCAAAGGAGAAGAACTCTTCACTGGAGTTGTCCCAATTC<br>TTGTTGAATTAGATGGTGATGTTAACGGCCACAAGTTCTCTGTCAGT<br>GGAGAGGGTGAAGGTGATGCAACATACGGAAAACTTACCCTGAAGT<br>TCATCTGCACTACTGGCAAACTGCCTGTTCCATGGCCAACACTAGTC<br>ACTACTCTGTGCTATGGTGTTCAATGCTTTTCAAGATACCCGGATCAT<br>ATGAAACGGCATGACTTTTTCAAGAGTGCCATGCCCGAAGGTTATGT<br>ACAGGAAAGGACCATCTTCTTCAAAGATGACGGCAACTACAAGACA<br>CGTGCTGAAGTCAAGTTTGAAGGTGATACCCTTGTTAATAGAATCGA<br>GTTAAAAGGTATTGACTTCAAGGAAGATGGCAACATTCTGGGACAC<br>AAATTGGAATACAACTATAACTCACACAATGTATACATCATGGCAGAC<br>AAACAAAAGAATGGAATCAAAGTGAACTTCAAGACCCGCCACAAC<br>ATTGAAGATGGAAGCGTTCAACTAGCAGACCATTATCAACAAAATAC<br>TCCAATTGGCGATGGCCCTGTCCTTTTACCAGACAACCATTACCTGTC<br>CACACAATCTGCCCTTTCGAAAGATCCCAACGAAAAGAGAGACCAC<br>ATGGTCCTTCTTGAGTTTGTAACAGCTGCTGGGATTACACATGGCAT<br>GGATGAACTGTACAACTGA |
| *SOX7* | ATGAAAAGGCCGGCGGCCACGAAAAAGGCCGGCCAGGCAAAAAAG<br>AAAAAGGGTTCTGGAGCTTCGCTGCTGGGAGCCTACCCTTGGCCCG<br>AGGGTCTCGAGTGCCCGGCCCTGGACGCCGAGCTGTCGGATGGACA<br>ATCGCCGCCGGCCGTCCCCCGGCCCCCGGGGGACAAGGGCTCCGAG<br>AGCCGTATCCGGCGGCCCATGAACGCCTTCATGGTTTGGGCCAAGG<br>ACGAGAGGAAACGGCTGGCAGTGCAGAACCCGGACCTGCACAACG<br>CCGAGCTCAGCAAGATGCTGGGAAAGTCGTGGAAGGCGCTGACGCT<br>GTCCCAGAAGAGGCCGTACGTGGACGAGGCGGAGCGGCTGCGCCT<br>GCAGCACATGCAGGACTACCCCAACTACAAGTACCGGCCGCGCAGG<br>AAGAAGCAGGCCAAGCGGCTGTGCAAGCGCGTGGACCCGGGCTTC<br>CTTCTGAGCTCCCTCTCCCGGGACCAGAACGCCCTGCCGGAGAAGA<br>GAAGCGGCAGCCGGGGGGCGCTGGGGGAGAAGGAGGACAGGGGT<br>GAGTACTCCCCCGGCACTGCCCTGCCCAGCCTCCGGGGCTGCTACC<br>ACGAGGGGCCGGCTGGTGGTGGCGGCGGCGGCACCCCGAGCAGTG<br>TGGACACGTACCCGTACGGGCTGCCCACACCTCCTGAAATGTCTCCC<br>CTGGACGTGCTGGAGCCGGAGCAGACCTTCTTCTCCTCCCCCTGCC<br>AGGAGGAGCATGGCCATCCCCGCCGCATCCCCCACCTGCCAGGGCA<br>CCCGTACTCACCGGAGTACGCCCCAAGCCCTCTCCACTGTAGCCACC<br>CCCTGGGCTCCCTGGCCCTTGGCCAGTCCCCCGGCGTCTCCATGATG<br>TCCCCTGTACCCGGCTGTCCCCCATCTCCTGCCTATTACTCCCCGGCC<br>ACCTACCACCCACTCCACTCCAACCTCCAAGCCCACCTGGGCCAGC<br>TTTCCCCGCCTCCTGAGCACCCTGGCTTCGACGCCCTGGATCAACTG<br>AGCCAGGTGGAACTCCTGGGGGACATGGATCGCAATGAATTCGACC<br>AGTATTTGAACACTCCTGGCCACCCAGACTCCGCCACAGGGGCCAT<br>GGCCCTCAGTGGGCATGTTCCGGTCTCCCAGGTGACACCAACGGGT |

| | |
|---|---|
| | CCCACAGAGACCAGCCTCATCTCCGTCCTGGCTGATGCCACGGCCA<br>CGTACTACAACAGCTACAGTGTGTCATGA |
| *S-gene* | ATGTTTGTTTTTCTTGTTTTATTGCCACTAGTCTCTAGTCAGTGTGTTA<br>ATCTTACAACCAGAACTCAATTACCCCCTGCATACACTAATTCTTTCA<br>CACGTGGTGTTTATTACCCTGACAAAGTTTTCAGATCCTCAGTTTTAC<br>ATTCAACTCAGGACTTGTTCTTACCTTTCTTTTCCAATGTTACTTGGT<br>TCCATGCTATACATGTCTCTGGGACCAATGGTACTAAGAGGTTTGATA<br>ACCCTGTCCTACCATTTAATGATGGTGTTTATTTTGCTTCCACTGAGA<br>AGTCTAACATAATAAGAGGCTGGATTTTTGGTACTACTTTAGATTCGA<br>AGACCCAGTCCCTACTTATTGTTAATAACGCTACTAATGTTGTTATTAA<br>AGTCTGTGAATTTCAATTTTGTAATGATCCATTTTTGGGTGTTTATTAC<br>CACAAAAACAACAAAAGTTGGATGGAAAGTGAGTTCAGAGTTTATT<br>CTAGTGCGAATAATTGCACTTTTGAATATGTCTCTCAGCCTTTTCTTAT<br>GGACCTTGAAGGAAAACAGGGTAATTTCAAAAATCTTAGGGAATTT<br>GTGTTTAAGAATATTGATGGTTATTTTAAAATATATTCTAAGCACACGC<br>CTATTAATTTAGTGCGTGATCTCCCTCAGGGTTTTTCGGCTTTAGAAC<br>CATTGGTAGATTTGCCAATAGGTATTAACATCACTAGGTTTCAAACTT<br>TACTTGCTTTACATAGAAGTTATTTGACTCCTGGTGATTCTTCTTCAG<br>GTTGGACAGCTGGTGCTGCAGCTTATTATGTGGGTTATCTTCAACCTA<br>GGACTTTTCTATTAAAATATAATGAAAATGGAACCATTACAGATGCTG<br>TAGACTGTGCACTTGACCCTCTCTCAGAAACAAAGTGTACGTTGAA<br>ATCCTTCACTGTAGAAAAAGGAATCTATCAAACTTCTAACTTTAGAG<br>TCCAACCAACAGAATCTATTGTTAGATTTCCTAATATTACAAACTTGT<br>GCCCTTTTGGTGAAGTTTTTAACGCCACCAGATTTGCATCTGTTTATG<br>CTTGGAACAGGAAGAGAATCAGCAACTGTGTTGCTGATTATTCTGTC<br>CTATATAATTCCGCATCATTTTCCACTTTTAAGTGTTATGGAGTGTCTC<br>CTACTAAATTAAATGATCTCTGCTTTACTAATGTCTATGCAGATTCATT<br>TGTAATTAGAGGTGATGAAGTCAGACAAATCGCTCCAGGGCAAACT<br>GGAAAGATTGCTGATTATAATTATAAATTACCAGATGATTTTACAGGC<br>TGCGTTATAGCTTGGAATTCTAACAATCTTGATTCTAAGGTTGGTGGT<br>AATTATAATTACCTGTATAGATTGTTTAGGAAGTCTAATCTCAAACCTT<br>TTGAGAGAGATATTTCAACTGAAATCTATCAGGCCGGTAGCACACCT<br>TGTAATGGTGTTGAAGGTTTTAATTGTTACTTTCCTTTACAATCATATG<br>GTTTCCAACCCACTAATGGTGTTGGTTACCAACCATACAGAGTAGTA<br>GTACTTTCTTTTGAACTTCTACATGCACCAGCAACTGTTTGTGGACCT<br>AAAAAGTCTACTAATTTGGTTAAAAACAAATGTGTCAATTTCAACTT<br>CAATGGTTTAACAGGCACAGGTGTTCTTACTGAGTCTAACAAAAAGT<br>TTCTGCCTTTCCAACAATTTGGCAGAGACATTGCTGACACTACTGAT<br>GCTGTCCGTGATCCACAGACACTTGAGATTCTTGACATTACACCATG<br>TTCTTTTGGTGGTGTCAGTGTTATAACACCAGGAACAAATACTTCTA<br>ACCAGGTTGCTGTTCTTTATCAGGATGTTAACTGCACAGAAGTCCCT<br>GTTGCTATTCATGCAGATCAACTTACTCCTACTTGGCGTGTTTATTCTA<br>CAGGTTCTAATGTTTTTCAAACACGTGCAGGCTGTTTAATAGGGGCT<br>GAACATGTCAACAACTCATATGAGTGTGACATACCCATTGGTGCAGG |

| | |
|---|---|
| | TATATGCGCTAGTTATCAGACTCAGACTAATTCTCCTCGGCGGGCACG<br>TAGTGTAGCTAGTCAATCCATCATTGCCTACACTATGTCACTTGGTGC<br>AGAAAATTCAGTTGCTTACTCTAATAACTCTATTGCCATACCCACAAA<br>TTTTACTATTAGTGTTACCACAGAAATTCTACCAGTGTCTATGACCAA<br>GACATCAGTAGATTGTACAATGTACATTTGTGGTGATTCAACTGAATG<br>CAGCAATCTTTTGTTGCAATATGGCAGTTTTTGTACACAATTAAACCG<br>TGCTTTAACTGGAATAGCTGTTGAACAAGACAAAAACACCCAAGAA<br>GTTTTTGCACAAGTCAAACAAATTTACAAAACACCACCAATTAAAG<br>ATTTTGGTGGTTTTAATTTTTCACAAATATTACCAGATCCATCAAAAC<br>CAAGCAAGAGGTCATTTATTGAAGATCTACTTTTCAACAAAGTGACA<br>CTTGCAGATGCTGGCTTCATCAAACAATATGGTGATTGCCTTGGTGAT<br>ATTGCTGCTAGAGACCTCATTTGTGCACAAAAGTTTAACGGCCTTAC<br>TGTTTTGCCACCTTTGCTCACAGATGAAATGATTGCTCAATACACTTC<br>TGCACTGTTAGCGGGTACAATCACTTCTGGTTGGACCTTTGGTGCAG<br>GTGCTGCATTACAAATACCATTTGCTATGCAAATGGCTTATAGGTTTA<br>ATGGTATTGGAGTTACACAGAATGTTCTCTATGAGAACCAAAAATTG<br>ATTGCCAACCAATTTAATAGTGCTATTGGCAAAATTCAAGACTCACTT<br>TCTTCCACAGCAAGTGCACTTGGAAAACTTCAAGATGTGGTCAACC<br>AAAATGCACAAGCTTTAAACACGCTTGTTAAACAACTTAGCTCCAAT<br>TTTGGTGCAATTTCAAGTGTTTTAAATGATATCCTTTCACGTCTTGAC<br>AAAGTTGAGGCTGAAGTGCAAATTGATAGGTTGATCACAGGCAGAC<br>TTCAAAGTTTGCAGACATATGTGACTCAACAATTAATTAGAGCTGCA<br>GAAATCAGAGCTTCTGCTAATCTTGCTGCTACTAAAATGTCAGAGTG<br>TGTACTTGGACAATCAAAAAGAGTTGATTTTTGTGGAAAGGGCTATC<br>ATCTTATGTCCTTCCCTCAGTCAGCACCTCATGGTGTAGTCTTCTTGC<br>ATGTGACTTATGTCCCTGCACAAGAAAAGAACTTCACAACTGCTCCT<br>GCCATTTGTCATGATGGAAAAGCACACTTTCCTCGTGAAGGTGTCTT<br>TGTTTCAAATGGCACACACTGGTTTGTAACACAAAGGAATTTTTATG<br>AACCACAAATCATTACTACAGACAACACATTTGTGTCTGGTAACTGT<br>GATGTTGTAATAGGAATTGTCAACAACACAGTTTATGATCCTTTGCAA<br>CCTGAATTAGACTCATTCAAGGAGGAGTTAGATAAATATTTTAAGAAT<br>CATACATCACCAGATGTTGATTTAGGTGACATCTCTGGCATTAATGCT<br>TCAGTTGTAAACATTCAAAAAGAAATTGACCGCCTCAATGAGGTTGC<br>CAAGAATTTAAATGAATCTCTCATCGATCTCCAAGAACTTGGAAAGT<br>ATGAGCAGTATATAAAATGGCCATGGTACATTTGGCTAGGTTTTATAG<br>CTGGCTTGATTGCCATAGTAATGGTGACAATTATGCTTTGCTGTATGA<br>CCAGTTGCTGTAGTTGTCTCAAGGGCTGTTGTTCTTGTGGATCCTGC<br>TGCAAATTTGATGAAGACGACTCTGAGCCAGTGCTCAAAGGAGTCA<br>AATTACATTACACATAA |
| *Cas9* | ATGAAAAGGCCGGCGGCCACGAAAAAGGCCGGCCAGGCAAAAAAG<br>AAAAAGGGTTCTGGAGATAAAAAGTATTCTATTGGTTTAGACATCGG<br>CACTAATTCCGTTGGATGGGCTGTCATAACCGATGAATACAAAGTAC<br>CTTCAAAGAAATTTAAGGTGTTGGGGAACACAGACCGTCATTCGATT<br>AAAAAGAATCTTATCGGTGCCCTCCTATTCGATAGTGGCGAAACGGC |

| | |
|---|---|
| | AGAGGCGACTCGCCTGAAACGAACCGCTCGGAGAAGGTATACACGT<br>CGCAAGAACCGAATATGTTACTTACAAGAAATTTTTAGCAATGAGAT<br>GGCCAAAGTTGACGATTCTTTCTTTCACCGTTTGGAAGAGTCCTTCC<br>TTGTCGAAGAGGACAAGAAACATGAACGGCACCCCATCTTTGGAAA<br>CATAGTAGATGAGGTGGCATATCATGAAAAGTACCCAACGATTTATCA<br>CCTCAGAAAAAAGCTAGTTGACTCAACTGATAAAGCGGACCTGAGG<br>TTAATCTACTTGGCTCTTGCCCATATGATAAAGTTCCGTGGGCACTTT<br>CTCATTGAGGGTGATCTAAATCCGGACAACTCGGATGTCGACAAACT<br>GTTCATCCAGTTAGTACAAACCTATAATCAGTTGTTTGAAGAGAACC<br>CTATAAATGCAAGTGGCGTGGATGCGAAGGCTATTCTTAGCGCCCGC<br>CTCTCTAAATCCCGACGGCTAGAAAACCTGATCGCACAATTACCCGG<br>AGAGAAGAAAAATGGGTTGTTCGGTAACCTTATAGCGCTCTCACTAG<br>GCCTGACACCAAATTTTAAGTCGAACTTCGACTTAGCTGAAGATGCC<br>AAATTGCAGCTTAGTAAGGACACGTACGATGACGATCTCGACAATCT<br>ACTGGCACAAATTGGAGATCAGTATGCGGACTTATTTTTGGCTGCCA<br>AAAACCTTAGCGATGCAATCCTCCTATCTGACATACTGAGAGTTAATA<br>CTGAGATTACCAAGGCGCCGTTATCCGCTTCAATGATCAAAAGGTAC<br>GATGAACATCACCAAGACTTGACACTTCTCAAGGCCCTAGTCCGTCA<br>GCAACTGCCTGAGAAATATAAGGAAATATTCTTTGATCAGTCGAAAA<br>ACGGGTACGCAGGTTATATTGACGGCGGAGCGAGTCAAGAGGAATT<br>CTACAAGTTTATCAAACCCATATTAGAGAAGATGGATGGGACGGAAG<br>AGTTGCTTGTAAAACTCAATCGCGAAGATCTACTGCGAAAGCAGCG<br>GACTTTCGACAACGGTAGCATTCCACATCAAATCCACTTAGGCGAAT<br>TGCATGCTATACTTAGAAGGCAGGAGGATTTTTATCCGTTCCTCAAAG<br>ACAATCGTGAAAAGATTGAGAAAATCCTAACCTTTCGCATACCTTAC<br>TATGTGGGACCCCTGGCCCGAGGGAACTCTCGGTTCGCATGGATGAC<br>AAGAAAGTCCGAAGAAACGATTACTCCCTGGAATTTTGAGGAAGTT<br>GTCGATAAAGGTGCGTCAGCTCAATCGTTCATCGAGAGGATGACCGC<br>CTTTGACAAGAATTTACCGAACGAAAAAGTATTGCCTAAGCACAGTT<br>TACTTTACGAGTATTTCACAGTGTACAATGAACTCACGAAAGTTAAG<br>TATGTCACTGAGGGCATGCGTAAACCCGCCTTTCTAAGCGGAGAACA<br>GAAGAAAGCAATAGTAGATCTGTTATTCAAGACCAACCGCAAAGTG<br>ACAGTTAAGCAATTGAAAGAGGACTACTTTAAGAAAATTGAATGCTT<br>CGATTCTGTCGAGATCTCCGGGGTAGAAGATCGATTTAATGCGTCAC<br>TTGGTACGTATCATGACCTCCTAAAGATAATTAAAGATAAGGACTTCC<br>TGGATAACGAAGAGAATGAAGATATCTTAGAAGATATAGTGTTGACT<br>CTTACCCTCTTTGAAGATCGGGAAATGATTGAGGAAAGACTAAAAA<br>CATACGCTCACCTGTTCGACGATAAGGTTATGAAACAGTTAAAGAGG<br>CGTCGCTATACGGGCTGGGGAGCCTTGTCGCGGAAACTTATCAACGG<br>GATAAGAGACAAGCAAAGTGGTAAAACTATTCTCGATTTTCTAAAGA<br>GCGACGGCTTCGCCAATAGGAACTTTATGGCCCTGATCCATGATGAC<br>TCTTTAACCTTCAAAGAGGATATACAAAAGGCACAGGTTTCCGGACA<br>AGGGGACTCATTGCACGAACATATTGCGAATCTTGCTGGTTCGCCAG<br>CCATCAAAAAGGGCATACTCCAGACAGTCAAAGTAGTGGATGAGCT |

| | |
|---|---|
| | AGTTAAGGTCATGGGACGTCACAAACCGGAAAACATTGTAATCGAG<br>ATGGCACGCGAAAATCAAACGACTCAGAAGGGGCAAAAAAACAGT<br>CGAGAGCGGATGAAGAGAATAGAAGAGGGTATTAAAGAACTGGGC<br>AGCCAGATCTTAAAGGAGCATCCTGTGGAAAATACCCAATTGCAGA<br>ACGAGAAACTTTACCTCTATTACCTACAAAATGGAAGGGACATGTAT<br>GTTGATCAGGAACTGGACATAAACCGTTTATCTGATTACGACGTCGA<br>TCACATTGTACCCCAATCCTTTTTGAAGGACGATTCAATCGACAATAA<br>AGTGCTTACACGCTCGGATAAGAACCGAGGGAAAAGTGACAATGTT<br>CCAAGCGAGGAAGTCGTAAAGAAAATGAAGAACTATTGGCGGCAGC<br>TCCTAAATGCGAAACTGATAACGCAAAGAAAGTTCGATAACTTAACT<br>AAAGCTGAGAGGGGTGGCTTGTCTGAACTTGACAAGGCCGGATTTA<br>TTAAACGTCAGCTCGTGGAAACCCGCGCCATCACAAAGCATGTTGC<br>GCAGATACTAGATTCCCGAATGAATACGAAATACGACGAGAACGATA<br>AGCTGATTCGGGAAGTCAAAGTAATCACTTTAAAGTCAAAATTGGTG<br>TCGGACTTCAGAAAGGATTTTCAATTCTATAAAGTTAGGGAGATAAA<br>TAACTACCACCATGCGCACGACGCTTATCTTAATGCCGTCGTAGGGA<br>CCGCACTCATTAAGAAATACCCGAAGCTAGAAAGTGAGTTTGTGTAT<br>GGTGATTACAAAGTTTATGACGTCCGTAAGATGATCGCGAAAAGCGA<br>ACAGGAGATAGGCAAGGCTACAGCCAAATACTTCTTTTATTCTAACA<br>TTATGAATTTCTTTAAGACGGAAATCACTCTGGCAAACGGAGAGATA<br>CGCAAACGACCTTTAATTGAAACCAATGGGGAGACAGGTGAAATCG<br>TATGGGATAAGGGCCGGGACTTCGCGACGGTGAGAAAAGTTTTGTC<br>CATGCCCCAAGTCAACATAGTAAAGAAAACTGAGGTGCAGACCGGA<br>GGGTTTTCAAAGGAATCGATTCTTCCAAAAAGGAATAGTGATAAGCT<br>CATCGCTCGTAAAAAGGACTGGGACCCGAAAAAGTACGGTGGCTTC<br>GATAGCCCTACAGTTGCCTATTCTGTCCTAGTAGTGGCAAAAGTTGA<br>GAAGGGAAAATCCAAGAAACTGAAGTCAGTCAAAGAATTATTGGGG<br>ATAACGATTATGGAGCGCTCGTCTTTTGAAAAGAACCCCATCGACTT<br>CCTTGAGGCGAAAGGTTACAAGGAAGTAAAAAAGGATCTCATAATT<br>AAACTACCAAAGTATAGTCTGTTTGAGTTAGAAAATGGCCGAAAAC<br>GGATGTTGGCTAGCGCCGGAGAGCTTCAAAAGGGGAACGAACTCGC<br>ACTACCGTCTAAATACGTGAATTTCCTGTATTTAGCGTCCCATTACGA<br>GAAGTTGAAAGGTTCACCTGAAGATAACGAACAGAAGCAACTTTTT<br>GTTGAGCAGCACAAACATTATCTCGACGAAATCATAGAGCAAATTTC<br>GGAATTCAGTAAGAGAGTCATCCTAGCTGATGCCAATCTGGACAAAG<br>TATTAAGCGCATACAACAAGCACAGGGATAAACCCATACGTGAGCAG<br>GCGGAAAATATTATCCATTTGTTTACTCTTACCAACCTCGGCGCTCCA<br>GCCGCATTCAAGTATTTTGACACAACGATAGATCGCAAACGATACAC<br>TTCTACCAAGGAGGTGCTAGACGCGACACTGATTCACCAATCCATCA<br>CGGGATTATATGAAACTCGGATAGATTTGTCACAGCTTGGGGGTGAC |
| 5´UTR | GGACAGATCGCCTGGAGACGCCATCCACGCTGTTTTGACCTCCATAG<br>AAGACACCGGGACCGATCCAGCCTCCGCGGCCGGGAACGGTGCATT<br>GGAACGCGGATTCCCCGTGCCAAGAGTGACTCACCGTCCTTGACAC<br>G |

| | |
|---|---|
| T1-F-primer | CCAGGGTTTTCCCAGTCACGACGTTGTAAAACGAC |
| T2-F-primer | GGTGTTGGCGGGTGTCGGGGCTGGCTTAACTATGC |
| T3-F-primer | GAAGCATTTATCAGGGTTATTGTCTCATGAGCGGA |
| *Gfp*-template-R | TCAGTTGTACAGTTCATCCATGCCATGTGTAATCCC |
| *SOX7*-template-R | TCATGACACACTGTAGCTGTTGTAGTACGTGGCCGT |
| *S-gene*-template-R | TTATGTGTAATGTAATTTGACTCCTTTGAGCACTG |
| *Cas9*-template-R | TCACCCCCAAGCTGTGACAAATCTATCCGAGTTTC |
| *Gfp*-template-CTGT-R | ACAGTTGTACAGTTCATCCATGCCATGTGTAATCCC |
| *Gfp*-template-CTGC-R | GCAGTTGTACAGTTCATCCATGCCATGTGTAATCCC |
| *Gfp*-template-CTGG-R | CCAGTTGTACAGTTCATCCATGCCATGTGTAATCCC |
| *Gfp*-template-AAAA-R | TTTTTCAGTTGTACAGTTCATCCATGCCATGTGTAATCCC |
| *Gfp*-template-TTTT-R | AAAATCAGTTGTACAGTTCATCCATGCCATGTGTAATCCC |
| *Gfp*-template-CCCC-R | GGGGTCAGTTGTACAGTTCATCCATGCCATGTGTAATCCC |
| *Gfp*-template-GGGG-R | CCCCTCAGTTGTACAGTTCATCCATGCCATGTGTAATCCC |
| *Gfp*-template-1,2,3,4T-R | AAAATTGTACAGTTCATCCATGCCATGTGTAATCCC |
| *Gfp*-template-1A-R | TAAATTGTACAGTTCATCCATGCCATGTGTAATCCC |
| *Gfp*-template-1C-R | GAAATTGTACAGTTCATCCATGCCATGTGTAATCCC |
| *Gfp*-template-1G-R | CAAATTGTACAGTTCATCCATGCCATGTGTAATCCC |
| *Gfp*-template-2A-R | ATAATTGTACAGTTCATCCATGCCATGTGTAATCCC |
| *Gfp*-template-2C-R | AGAATTGTACAGTTCATCCATGCCATGTGTAATCCC |
| *Gfp*-template-2G-R | ACAATTGTACAGTTCATCCATGCCATGTGTAATCCC |
| *Gfp*-template-3A-R | AATATTGTACAGTTCATCCATGCCATGTGTAATCCC |
| *Gfp*-template-3C-R | AAGATTGTACAGTTCATCCATGCCATGTGTAATCCC |

| | |
|---|---|
| *Gfp*-template-3G-R | AACATTGTACAGTTCATCCATGCCATGTGTAATCCC |
| *Gfp*-template-4A-R | AAATTTGTACAGTTCATCCATGCCATGTGTAATCCC |
| *Gfp*-template-4C-R | AAAGTTGTACAGTTCATCCATGCCATGTGTAATCCC |
| *Gfp*-template-4G-R | AAACTTGTACAGTTCATCCATGCCATGTGTAATCCC |
| DNA without promoter | TTTAAAGGTGTCCGATGGATCTTCAGAGATCTTCTTTGGG |
| DNA with T7 promoter | TAATACGACTCACTATATTTAAAGGTGTCCGATGGATCTTCAGAGATCTTCTTTGGG |
| 53-bp template | CTAATACGACTCACTATAGGGAGACCCTCGAGGACAGATCGCCTGGAGACGCC |
| RACE linker | GAUAAAAAGUAUUCUAUUGGUUUAGACAUCGGCACUAAUUCCGUUGGAUGGGCUGUCAUAACCGAUGAAUACAAAGUACCUUCAAAGAAAUUUAAGGUGUUGGGGAACACAGACCGUCAUUCGAUUAAAAAGAAUCUUAUCGGUGCCCUCCUAUUCGAUAGUGGCGAAACGGCAGAGGCGACUCGCCUGAAACGAACCGC |
| 3’RACE-gene-F | GCCACCTATACTTTCGCCAGCTGGCGTAATAGCGA |
| 3’RACE-gene-R | CAGAGGCGACTCGCCTGAAACGAACCGCTATAGGT |
| 3’RACE-plasmid-F | CGAACCGCTATAGGTGGCCCAATTAAGAATTCACT |
| 3’RACE-plasmid-R | CTGGCGAAAGTATAGGTGGCCCAATTAAAAGCTTG |
| 5’RACE-gene-F | ACCTATAGATAAAAAGTATTCTATTGGTTTAGAC |
| 5’RACE-gene-R | ACCTATAATGGAAGCGTTCAACTAGCAGACCATTATCA |
| 5’RACE-plasmid-F | CGCTTCCATTATAGGTGGCCCAATTAAGAATTCACTGG |
| ’RACE-plasmid-R | CTTTTTATCTATAGGTGGCCCAATTAAAAGCTTGGC |

**Table S2**. 3´RACE results of antisense RNA.

| 3´RACE | | |
|---|---|---|
| | **Sequences** | **Reads/Total** |
| 3´-terminal sequence of DNA template | ACACCCGCCAACACC | / |
| 3´RACE results of antisense RNA | ACACCCGCCAACACC | 1/10 |
| | ACACCCGCCAA - - - - | 1/10 |
| | ACACCCGCCAA - - - - - | 1/10 |
| | ACACCCGCCAACACCC | 6/10 |
| | ACACCCGCCAACACCCG | 1/10 |
